# Endogenous μ-opioid – neuropeptide Y Y1 receptor synergy silences chronic postoperative pain

**DOI:** 10.1101/2022.07.05.498802

**Authors:** Tyler S. Nelson, Diogo F. S. Santos, Pranav Prasoon, Margaret Gralinski, Heather N. Allen, Bradley K. Taylor

## Abstract

Tissue injury creates a delicate balance between latent pain sensitization (LS) and compensatory endogenous analgesia. Inhibitory G protein-coupled receptor (GPCR) interactions that oppose LS, including μ-opioid receptor (MOR) and neuropeptide Y Y1 receptor (Y1R) activity, persist in the spinal cord dorsal horn (DH) for months, even after the resolution of normal pain thresholds. Here, we demonstrate that following recovery from surgical incision, a potent endogenous analgesic synergy between MOR and Y1R activity persists within DH interneurons to reduce the intensity and duration of latent postoperative hyperalgesia and ongoing pain. Failure of such endogenous GPCR signaling to maintain LS in remission may underlie the transition from acute to chronic pain states.

## Introduction

Chronic postsurgical pain (CPSP) is a significant healthcare burden that afflicts millions of patients each year (Richebé et al., 2018; Thapa and Euasobhon, 2018). Despite this high prevalence, the biological mechanisms that underlie the transition from acute pain to CPSP remain poorly understood (Glare et al., 2019; Kehlet et al., 2006). The dorsal horn of the spinal cord (DH) processes somatosensory information and is a key driver of pathological pain states (Todd, 2010). Tissue injury sensitizes pro-nociceptive neurons in the DH, contributing to allodynia and hyperalgesia (Jensen and Finnerup, 2014; Kuner, 2010; Latremoliere and Woolf, 2009). However, accumulating evidence from human and animal studies suggest that after tissue injury-induced hyperalgesia resolves, sensitization in the DH persists within a long-lasting silent state of remission, termed “latent sensitization” (LS) (Gerum and Simonin, 2021; Taylor and Corder, 2014).

Following tissue injury and the subsequent resolution of hyperalgesia, intrathecal administration (i.t.) of selective antagonists at inhibitory Gα_i/o_ G protein-coupled receptors (GPCRs), including μ-opioid receptors (MOR), kappa opioid receptors (KOR), neuropeptide Y Y1 receptors (Y1R), or several other receptors, unmask LS and reinstate hyperalgesia (Basu et al., 2021; Corder et al., 2013; Fu et al., 2019, 2020; Solway et al., 2011; Walwyn et al., 2016). Remarkably, each antagonist was sufficient to produce a complete, not partial, reinstatement of hyperalgesia. This suggested to us that GPCRs interact in a complex manner, not just additively, to maintain LS in remission. Indeed, individual cells express many GPCRs whose intracellular second messengers can interact to co-alter signaling (Cordeaux and Hill, 2002; Gupte et al., 2017; Hur and Kim, 2002; Selbie and Hill, 1998). For example, different GPCRs can activate the same G proteins (Alt et al., 2002; Yao et al., 2006). Thus, coincidental activation of second messenger pathways by co-activation of multiple GPCRs can elicit supra-additive (synergistic) amplification of the responses and produce a greater than additive leftward shift in the response curve (Aira et al., 2014; Bourne and Nicoll, 1993; Horioka et al., 2021; Philip et al., 2010).

Examples of spinal analgesic synergy between Gα_i/o_ GPCR agonists exist in the pharmacology literature, including mu and kappa-selective or mu and delta-selective opiates (Schuster et al., 2015; Sutters et al., 1990), opiates and cannabinoids (Cichewicz, 2004; Grenald et al., 2017; Kazantzis et al., 2016), and opiates and α2-adrenergic receptor agonists (Chabot-Doré et al., 2015; Overland et al., 2009; Stone et al., 1997). The aim of this study is to test the hypothesis that surgical incision produces a tonic and long-lasting synergistic dependence on MOR and Y1R endogenous signaling to oppose the development of CPSP.

## Results

### MOR and Y1R are co-expressed in DRG and DH

Synergistic interactions between MOR and Y1R may be mediated by either 1) intracellular mechanisms in which receptors located on the same cell produce interactions at the level of intracellular signaling cascades, or 2) via intercellular mechanisms which involve coincident inhibition of two neurons in series in the same anatomical pathway or a retrograde feedback mechanism (Chabot-Doré et al., 2015). First, we examined MOR and Y1R localization using fluorescence *in situ* hybridization for *Oprm1* and *Npy1r* and we found colocalization in cells in the lumbar dorsal root ganglion (DRG) (**Fig. 1A-B**) and DH (**Fig. 1C-D**). Thus, MOR and Y1R intracellular cross-talk in neurons in both DRG and DH is plausible to produce synergistic intracellular signaling.

**Fig 1:**
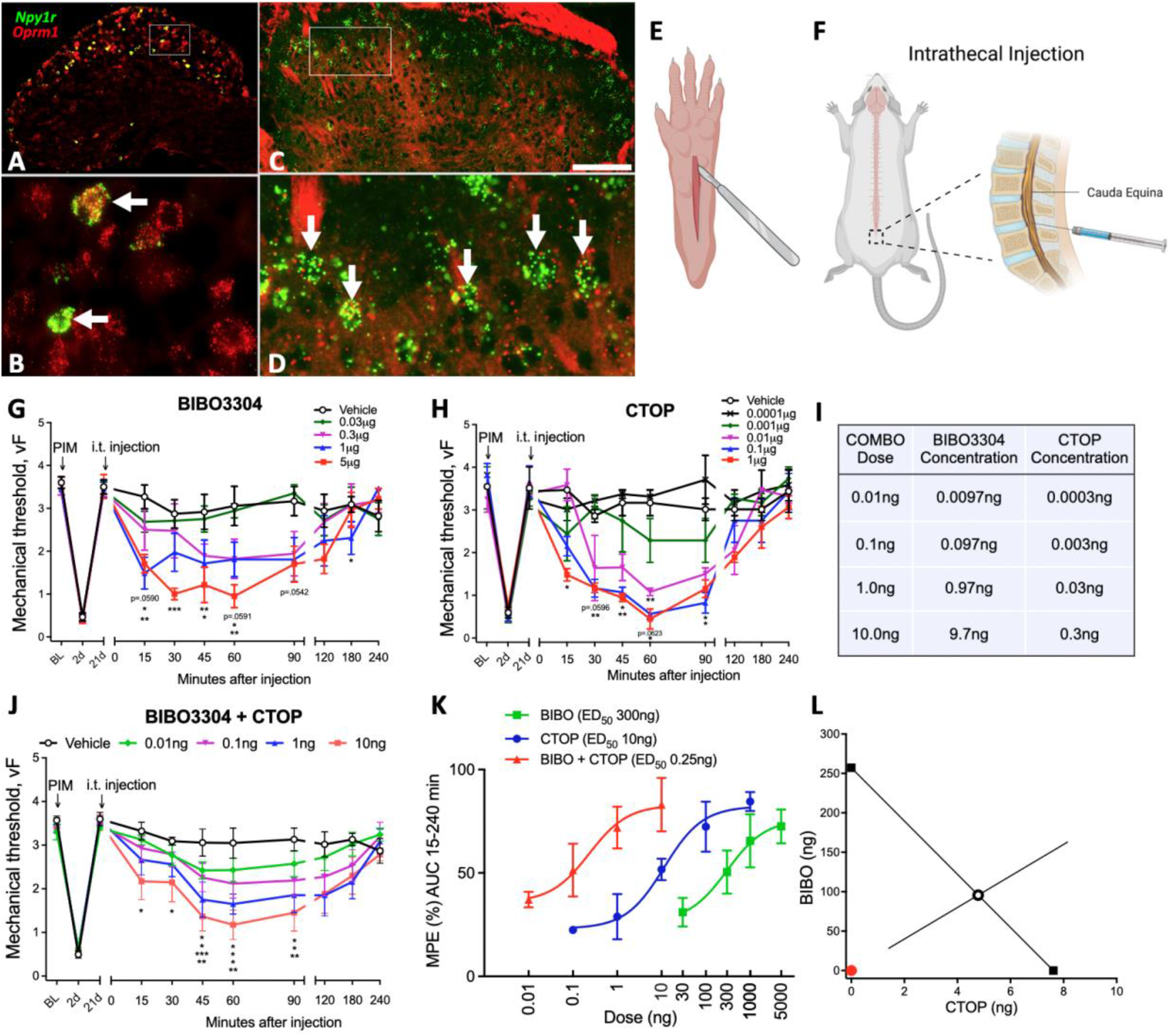
Y1R and MOR synergistically oppose LS. Fluorescence *in situ* hybridization demonstrating colocalization of *Oprm1* and *Npy1r* in the same cells in both lumbar DRG (**A-B**) and DH (**C-D**). **B** and **D** represent zoomed in box insets for **A** and **C**, respectively. Schematic depictions of plantar incision (**E**) and intrathecal injections into the mouse to target the spinal cord (**F**). Dose-response time courses of reinstatement of hyperalgesia after intrathecal (i.t.) administration of BIBO3304 (**G**), CTOP (**H**), or BIBO3304 and CTOP in combination (**I-J**) (n=3-8/group). Dose response effects of antagonist-induced reinstatement. MPE: maximum possible effect. AUC: area under curve (**K**). Isobolographic analysis of interaction between BIBO3304 and CTOP and red dot indicating interaction index of 0.02, a measure of drug synergism by which a value < 1.0 is determined to be synergistic (**L**). Data presented as mean ± SEM. Significance determined with two-way RM ANOVAs followed by post hoc if applicable with *P<0.05, **P<0.01, and ***P<0.001.

### MOR and Y1R signaling work synergistically to oppose CPSP

Next, we performed plantar incision of the hindpaw (PIM), a model of postoperative pain (**Fig. 1E**) that produces robust mechanical hyperalgesia that resolves within 21 days. Following resolution of hyperalgesia, i.t. administration (**Fig. 1F**) of a MOR antagonist (CTOP) or a Y1R antagonist (BIBO3304) dose-dependently reinstated mechanical hypersensitivity with an ED_50_ of 260ng and 8ng, respectively (**Fig. 1G-H**). Thus, CTOP exhibits a 30-fold difference in potency compared to BIBO3304, and we assessed synergistic interactions with a fixed ratio (30:1) isobologram method (Tallarida, 2016, 1992) (**Fig. 1I**). BIBO3304 and CTOP combination (BIBO:CTOP) reinstated mechanical hypersensitivity with robust effects at even a remarkably low 100pg dose (**Fig. 1J**). BIBO:CTOP produced a large leftward shift in the dose-response curve as compared to either BIBO3304 or CTOP administered alone (**Fig. 1K**), and the isobolographic analysis revealed a synergistic interaction (**Fig. 1L**). Additionally, BIBO:CTOP (10ng, i.t.) dramatically increased the light touch-evoked expression of phosphorylated extracellular signal-regulated kinase (pERK) in superficial DH neurons, a proxy for neuronal activation (**Fig. 2A-E**), and produced a robust conditioned place aversion in mice with plantar but not sham incision (**Fig. 2F-G**).

**Fig 2:**
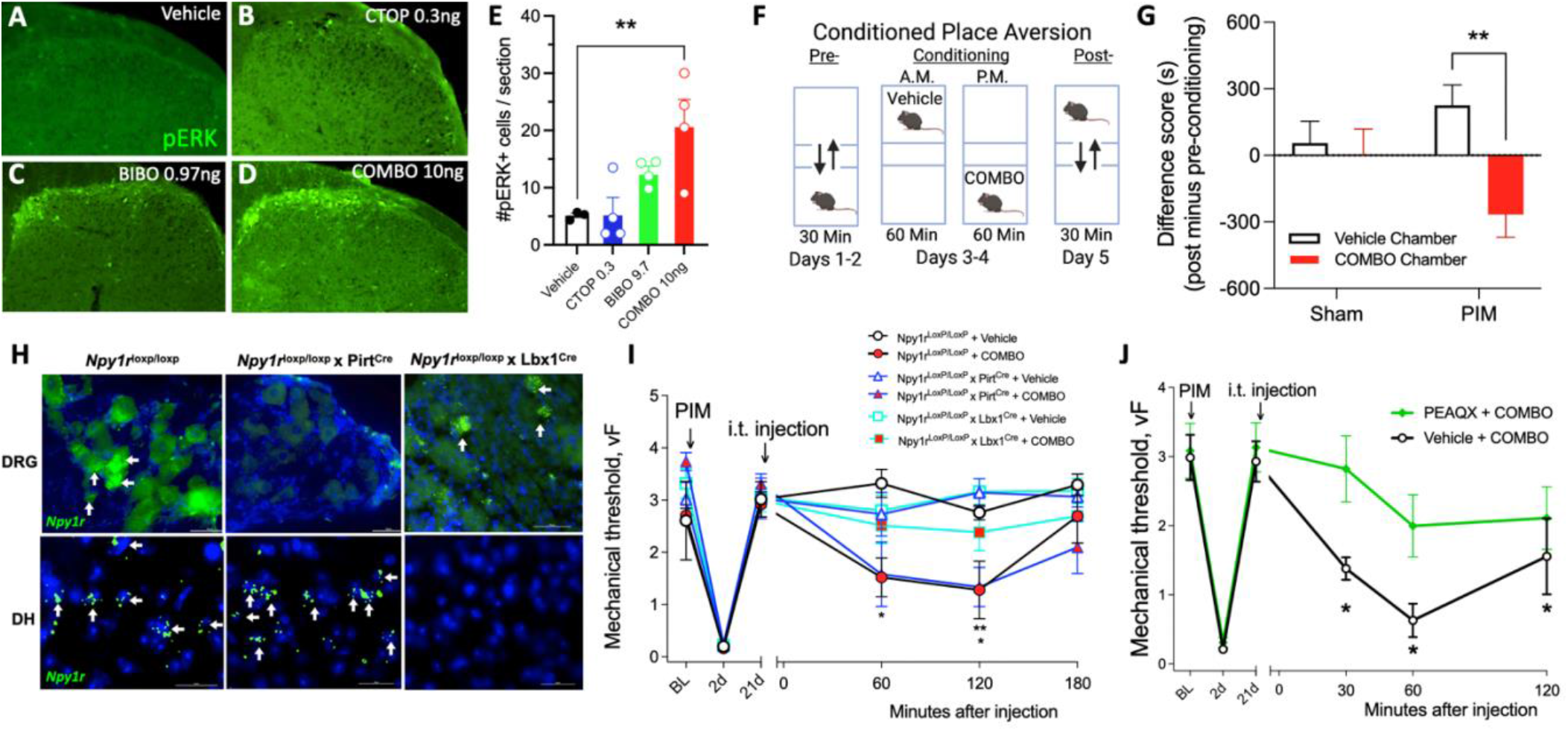
DH but not DRG Y1R and MOR synergy opposes a GluN2-driven LS. Representative images (**A-D**) and DH laminae I-II quantification of light touch-evoked pERK after intrathecal drug administration (**E**) (n=3-4). Experimental timeline (**F**) and quantification for conditioned place aversion testing (**G**) (n=11-12/group). Fluorescence *in situ* hybridization demonstrating loss of *Npy1r* expression in the DRG of Npy1r^loxP/loxP^ × Pirt^Cre^ mice and SC of Npy1r^loxP/loxP^ × Lbx1^Cre^ mice (**H**). BIBO:CTOP reinstated PIM-induced mechanical hyperalgesia in Npy1r^loxP/loxP^ and Npy1r^loxP/loxP^ × Pirt^Cre^ mice but not in Npy1r^loxP/loxP^ × Lbx1^Cre^ mice (**I**) (n=6-9/group). GluN2a NMDAR subtype antagonist, PEAQX (100ng, i.t.) prevented BIBO:CTOP-induced reinstatement of mechanical hyperalgesia (**J**) (n=7-8). Data presented as mean ± SEM. Significance determined using three- (H) or two-way RM ANOVAs (G, I) followed by post hoc if applicable with *P<0.05, and **P<0.01.

### MOR and Y1R signaling within DH rather than DRG neurons works synergistically to oppose LS

Intrathecally administered chemicals can engage both DH and DRG neurons. To resolve the specific site of action for BIBO:CTOP, we crossed Npy1r^loxP/loxP^ mice with either Pirt^Cre^ or Lbx1^Cre^ mice to conditionally knockout *Npy1r* in the DRG or DH, respectively (**Fig. 2H**). BIBO3304:CTOP (10ng, i.t.) reinstated PIM-induced mechanical hypersensitivity in both control (Npy1r^loxP/loxP^) and DRG conditional knockout mice (Npy1r^loxP/loxP^ × Pirt^Cre^), but not in DH conditional knockout mice (Npy1r^loxP/loxP^ × Lbx1^Cre^) (**Fig. 2I**). These data suggest that MOR and Y1R signal within DH neurons, rather than DRG neurons, to synergistically oppose LS and maintain postoperative pain in remission.

### Spinal LS is dependent on GluN2A NMDA receptors

Previously, we demonstrated that a N-methyl-D-aspartate receptor (NMDAR) blocker, MK-801 (dizocilipine), prevented either Y1R (Fu et al., 2019) or MOR (Corder et al., 2013) antagonist-induced reinstatement of peripheral inflammatory pain. Thus, we hypothesized that NMDARs, in particular the GluN2A subunit, also mediate the LS that is unmasked by BIBO:CTOP. To test this, we co-administered the GluN2A-preferring NMDAR antagonist, PEAQX (100ng, i.t.), with BIBO:CTOP (10ng, i.t.). PEAQX, but not vehicle, abolished the BIBO:CTOP-induced reinstatement of mechanical hypersensitivity (**Fig. 2J**). These data suggest that MOR and Y1R signaling synergistically opposes a GluN2A-mediated latent postoperative pain sensitization and thus maintains postoperative pain in remission (**Fig. 3**).

**Fig 3:**
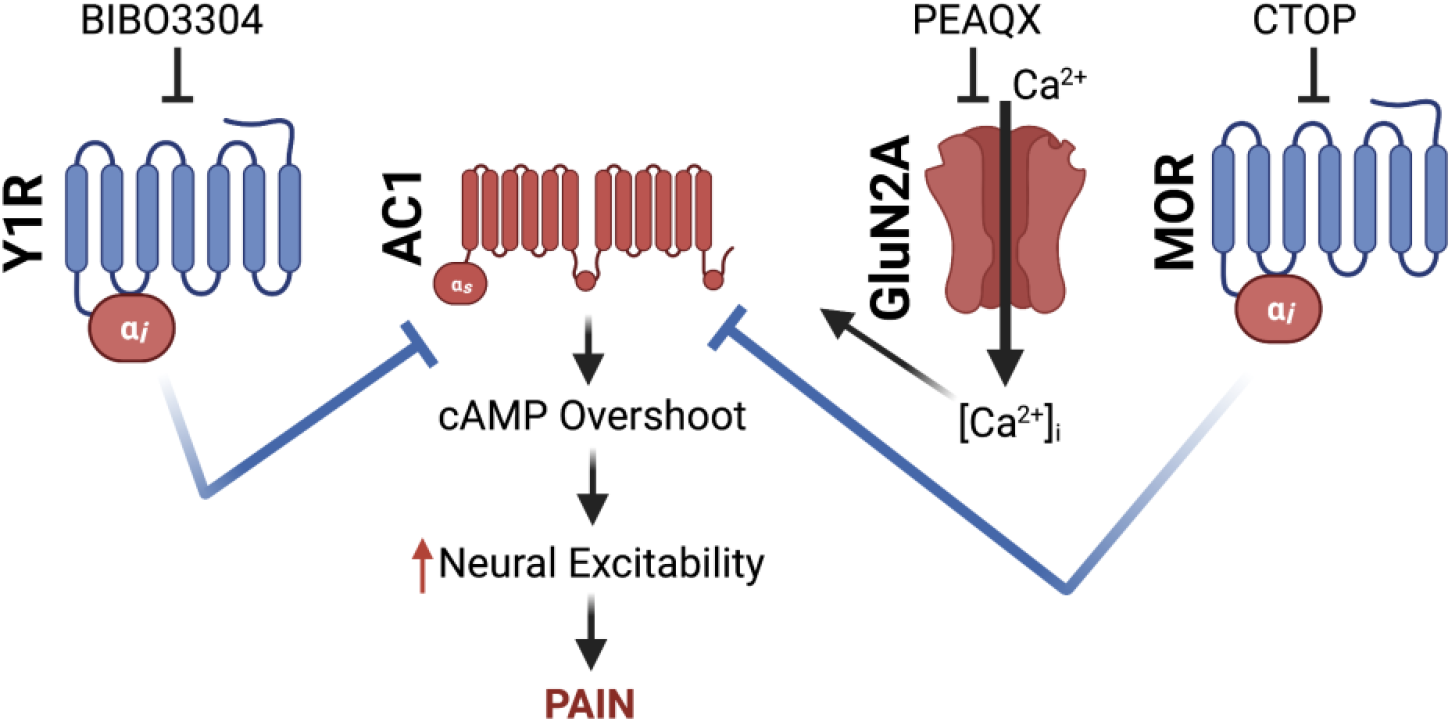
Proposed schematic of cellular pathways involved in endogenous NPY and opioid synergistic pain inhibition. We propose that following the resolution of injury, endogenous anti-nociceptive peptides (e.g. enkephalins, endorphine, NPY) interact with MOR and Y1R in a synergistic manner to maintain LS in remission. However, this long-lasting Gα_i/o_-coupled GPCR activity produces heterologous sensitization of adenylyl cyclase 1 (AC1). We hypothesize that both MOR and Y1R share a common pool of AC1, thus, potent activation or blockade of either MOR or Y1R can prevent or produce a cAMP overshoot and the reinstatement of hyperalgesia, respectively. This idea largely is proposed from the work of (Levitt et al., 2011).

## Discussion

Cells must integrate multiple signals from an array of receptors at any given moment. One of the most fundamental and evolutionarily conserved signaling mechanisms is GPCR activation, which is classically viewed as a compartmentalized cellular event in which a ligand binds a receptor to activate a specific signaling pathway distinct from other GPCRs. (Gurevich and Gurevich, 2020; Hur and Kim, 2002; Wang et al., 2018). However, researchers are uncovering examples of how GPCRs and their intracellular second messengers might interact within a cell to supra-additively co-alter signaling (Cordeaux and Hill, 2002; Gupte et al., 2017; Horioka et al., 2021; Hur and Kim, 2002; Overland et al., 2009; Schuster et al., 2015; Selbie and Hill, 1998). For the first time, we report an endogenous analgesic synergy between MOR and Y1R signaling that persists beyond the resolution of hyperalgesia and injury to maintain CPSP in remission. Modest failure in either Y1R or MOR compensatory signaling may underlie the physiological vulnerability to remission and the development of CPSP.

MOR and Y1R are *Pertussis* toxin-sensitive Gα_i/o_-coupled GPCRs; thus, upon initial receptor activation, the Gα_i/o_ subunit potently inhibits adenylyl cyclase to reduce the production of cAMP. The free Gβγ counterpart acts as a signaling molecule to activate downstream signaling pathways that include activation of G protein-coupled inwardly-rectifying K^+^ channels (GIRKs) and inhibition of voltage-gated Ca^2+^ channels to reduce the excitability of neurons (Yudin and Rohacs, 2018). Paradoxically, prolonged activation of Gα_i/o_ GPCRs enhances the activity of adenylyl cyclase and markedly increases cAMP production. This cellular phenomenon is referred to as heterologous sensitization (otherwise referred to as supersensitization, cAMP overshoot, cAMP superactivation) and is readily apparent upon removal of the agonist (Brust et al., 2015; Sharma et al., 1975; Watts, 2002). Interestingly, blockade of MOR constitutive activity in the setting of LS produces heterologous sensitization of adenylyl cyclase 1 (AC1) (Corder et al., 2013). This likely occurs for spinal Y1Rs as well, as the endogenous ligand, NPY, also produces heterologous sensitization (Drakulich et al., 2003), and Y1R antagonism-induced reinstatement of pain-like behavior is lost in AC1 knockout mice (Fu et al., 2019, 2020). Our current results suggest that endogenous anti-nociceptive peptides (e.g. enkephalins, endorphine, NPY) interact with MOR and Y1R in a synergistic manner to maintain LS in remission. Gα_i/o_-coupled GPCRs share a common pool of adenylyl cyclase, thus, when one Gα_i/o_-coupled GPCR produces heterologous sensitization, administration of a different Gα_i/o_ GPCR agonist can prevent subsequent cAMP overshoot (Levitt et al., 2011). As schematized in **Figure 3**, we suggest that endogenous MOR and Y1R activity synergistically inhibit AC1 while counter adaptively also producing a heterologous sensitization of AC1. Antagonism of the synergistically interacting, LS-inhibiting, Gα_i/o_-coupled GPCRs is therefore sufficient to evoke a cAMP overshoot and unmask LS to produce a complete reinstatement of hyperalgesia.

The molecular mechanisms underlying the endogenous MOR and Y1R synergy remain unknown, but several possible mechanisms exist. First, MOR and Y1R may form receptor-receptor interactions, such as the formation of heterodimers (Cordeaux and Hill, 2002; Selbie and Hill, 1998). The formation of heterodimers or oligomerization between GPCR receptors can markedly potentiate signal transduction (Jordan and Devi, 1999; Levac et al., 2002). Second, MOR and Y1R may undergo signal transduction interactions. The assumption is that Y1R and MOR coexist on neurons and share a common pool of G proteins; therefore, activation of one receptor may cause redistribution of its G proteins and increase the sensitivity of the other receptor. For example, binding of an endogenous ligand, such as NPY to Y1R, may shift the affinity of endogenous ligand binding to the separate GPCR MOR (Djellas et al., 2000). Additionally, both MOR and Y1R may synergistically work through downstream effectors. The free Gβγ released from the agonist-induced dissociation of both the MOR and Y1R Gi heterotrimers may co-activate protein kinase C (PKC), phospholipase C (PLC) (Overland et al., 2009; Yao et al., 2003), or protein kinase C epsilon (PKCε) (Schuster et al., 2015) to synergistically oppose LS. Third, peptide hormones like NPY can modulate neurotransmission by recruiting other GPCRs from the interior of the cell to the cell membrane (Holtbäck et al., 1999). Coincident activation of Y1R and MOR may allow recruitment of MORs to the cell membrane and a sensitization of MOR signal transduction (Achour et al., 2008; Cahill et al., 2007; Holtbäck et al., 1999). Future experiments should further evaluate how Y1R and MOR interact mechanistically to promote endogenous synergy.

The current study establishes the existence of supra-additive endogenous MOR and Y1R signaling in the spinal cord DH that maintains LS in remission. Further, we provide a strong basis for future investigations of the mechanisms involved in MOR-Y1R endogenous synergistic signaling and the cellular subpopulations in the DH that drive LS.

## Materials and Methods

### Animals

Adult C57Bl/6NCrl (Charles River, #027), Npy1r^loxP/loxP^ (courtesy of Herbert Herzog, (Howell et al., 2003)), Pirt^Cre^ (courtesy of Xinzhong Dong, (Kim et al., 2008)), and Lbx1^Cre^ (courtesy of Carmen Birchmeier, (Sieber et al., 2007)) mice were group housed, provided access to food and water *ad libitum*, and maintained on a 12:12 hour light:dark cycle (lights on at 7:00am) in temperature and humidity controlled rooms. Male and female mice were used in all experiments. No significant sex differences were observed. All experiments were carried out in accordance with guidelines from the International Association for the Study of Pain and with the approval of the Institutional Animal Care and Use Committees of the University of Pittsburgh.

### Drugs

| Chemical | Source | Dilutant Information |
| --- | --- | --- |
| BIBO 3304 trifluoroacetate | TOCRIS – Cat:2412 | Diluted in a vehicle solution of ETOH, castor oil and saline<br>in a 1:1:8 ratio |
| CTOP | TOCRIS – Cat:1578 | Diluted in saline |
| PEAQX tetrasodium salt | TOCRIS – Cat:5018 | Diluted in saline |

### Intrathecal injections

As previously described (Corder et al., 2013; Fu et al., 2019; Solway et al., 2011), the mouse was lightly restrained in a towel and a 30G ½ inch needle attached to a 25-μl Hamilton microsyringe was inserted into the subarachnoid space between the L5/L6 vertebrae at an angle of 30–45° to the horizontal plane. The needle was advanced until a reflexive tail flick was observed, at which time 5 μl of drug or vehicle was slowly administered. The needle was held in place for 10 seconds, withdrawn, and then the mouse was returned to its testing chamber.

### Synergistic Interaction/Isobologram Analysis

Drug interactions were evaluated by a statistical method known as isobolographic analysis in which the actual potency of two drugs in combination is compared to that predicted in the absence of an interaction (Tallarida, 2002). Isobolograms were constructed using the values obtained at the concentrations of the compounds administered alone and in combination that produced 50% of the possible maximum antinociceptive effect (ED_50_) in the Von Frey test. The theoretical dose required for a purely additive interaction (Z_add_) with the S.E.M. for each combination at a 1:1 ratio was computed from the ED_50_ values of the single drugs as previously described (Tallarida, 1992). The area under the curve (AUC) of mechanical withdrawal testing was calculated using the trapezoidal rule (Tallarida et al., 1989). Concentration-dependent curves for each of the tested compounds were established according to the percentage of antinociceptive effect that was calculated from the AUCs. The antinociceptive effect (%) was obtained from the AUCs of the different treatments relative to the AUC for vehicle. Isobolographics were generated using JFlashCalc (http://www.u.arizona.edu/~michaelo/jflashcalc.html) and reconstructed in GraphPad Prism 9.

The interaction index was calculated as follows:

A = ED_50_ from drug A alone (CTOP)
B = ED_50_ from drug B alone (BIBO 3304)
“a” and “b” = combination doses from the respective drugs based on the ED_50_ of the combination

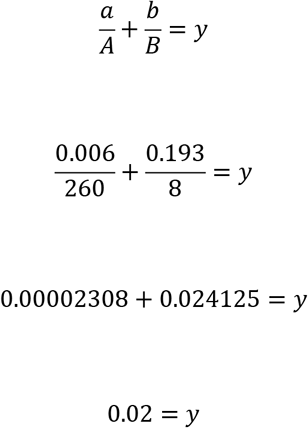 *Note that γ = 1 is additive, γ < 1 is supra-additive (synergistic), and γ >1 is sub-additive (antagonistic)

### Plantar Incision Model

Post-operative hyperalgesia was induced by longitudinal incision of the plantaris muscle as previously described (Basu et al., 2021; Pogatzki and Raja, 2003). Following antisepsis of the left hind paw with Chlorascrub® and 70% ethanol, a #11 scalpel blade was used to make a 5mm incision through the skin and fascia, beginning 2mm from the proximal edge of the heel and extending towards the digits. The underlying muscle was raised with a curved forceps, extended 4 mm, and then incised longitudinally with the #11 scalpel blade, all while leaving the origin and insertion of the muscle intact. The overlying skin was closed with synthetic 5-0 sutures (PDS*II, Ethicon). Surgery was typically completed within 5-10min. Surgeries were conducted under isoflurane anesthesia (5% induction followed by 1.5% - 2.0% maintenance). After suturing of the skin, triple antibiotic ointment (Neosporin, Johnson and Johnson) was applied to the surgical area. The sutures were removed 10 days after surgery.

### Behavioral Testing

#### Mechanical hypersensitivity

Sensitivity to a non-noxious mechanical stimulus was tested with an incremental series of 8 von Frey monofilaments of logarithmic stiffness (Stoelting, Wood Dale, IL) that ranged in gram force from 0.008g to 6g. The stimulation was applied lateral to the suture line. Filaments were applied to the skin with a slight bending of the filament for a maximum of 5 seconds. A clear withdrawal of the paw from the application of the stimulus was recorded as a positive response. The 50% withdrawal threshold was determined using the up-down method (Chaplan et al., 1994). Before commencement of each von Frey session, we acclimated the animals within individual Plexiglas boxes placed on the top of a stainless-steel mesh platform for 45 min.

#### Conditioned place aversion

A two-day conditioning protocol using a biased chamber assignment was used for conditioned place aversion (CPA). On the acclimation day (Day 0), mice had free access to explore all chambers of a 3-chamber conditioned place testing apparatus (side chambers: 170 x 150 mm; center chamber: 70 × 150 mm; height: 200 mm; San Diego Instruments) for 30 mins. Mice were able to discriminate between chambers using visual (vertical versus horizontal black-and-white striped walls) and sensory (rough versus smooth textured floor) cues. For pre-conditioning (Days 1 and 2), mice were again allowed to freely explore for 30 mins during which their position was recorded via a 4 × 16 infra-red photobeam array and associated software (San Diego Instruments). For conditioning (Days 3-4), each mouse’s non-preferred chamber was paired with a vehicle i.t. injection and the preferred chamber with a BIBO:CTOP combo i.t. injection. Each morning mice received an i.t. vehicle injection, were returned to their home cage for 5 min (to disassociate the injection with the chamber), and were then placed in the designated side chamber for 60 min. 6 hours later, mice received BIBO:CTOP combo (10ng, i.t.), were returned to their home cage for 5 min, and were placed into the BIBO:CTOP combo-designated chamber for 60 min. On test day (Day 5), mice could freely explore all chambers and their position was recorded as during pre-conditioning for 30 min. Difference scores were calculated as the time spent in the chamber on test day minus the time spent in the same chamber during pre-conditioning.

#### Touch-evoked phosphorylated extracellular signal-regulated kinase (pERK)

pERK was evoked by touch stimulation as previously described (Fu et al., 2019). 21 days after PIM, mice received either i.t. injections of vehicle, BIBO 3304 9.7ng, CTOP 0.3ng, or COMBO 10ng (BIBO 9.7ng + CTOP 0.3ng). One hour later, mice were lightly anesthetized with isoflurane (1.5%), and the ventral surface of the ipsilateral hindpaw was mechanically stimulated with a gentle 3-s stroke with a cotton swab from heel to toe. This was repeated every 5s for 5min. After an additional 5 min pause, mice were more deeply anesthetized with isoflurane and transcardially perfused with ice cold 0.01M phosphate buffered saline (PBS, Fischer Scientific), followed by 10% phosphate formalin buffer. Lumbar spinal cords were harvested and post-fixed in the same fixative overnight at 4 °C and then cryoprotected with 30% sucrose until total submersion (1–3 days).

#### Immunohistochemistry

Transverse spinal cord sections (30 μm) from L3-L5 were cut on a sliding microtome (Leica, SM, 2000R). A series of sections, each 240 μm apart, were washed in 0.01M PBS, blocked in 3% normal serum (goat; Gemini Bioproducts) containing 0.3% Triton X-100 (Sigma Aldrich) in 0.01M PBS for 1 h, and then incubated with primary rabbit antibody anti-phosphorylated-ERK1/2 antiserum (1:1000, Cell Signaling) at 4 °C for 24h on a shaker. The following day, sections were again washed in 0.01M PBS and incubated for 1 h at room temperature with the secondary conjugated antibody (1:1000, Invitrogen: goat anti-rabbit Alexa Fluor 488). The sections were washed in 0.01M phosphate buffer, mounted and coverslipped with VECTASHIELD HardSet Antifade Mounting Medium with DAPI. At least six good quality sections from segment L4 were selected from each subject for microscopy.

#### Fluorescence in situ Hybridization (FISH) (RNAscope)

Mice were transcardially perfused with ice cold 1× PBS followed by 10% buffered formalin and spinal cords and DRGs were extracted via blunt dissection, postfixed in 10% formalin (2-4 hrs), and then placed in 30% sucrose at 4°C until the tissue sank (~48-72 hrs). 20μm thick L3-L4 floating spinal cord sections were obtained on a vibrating microtome, and 12μm thick L3-L4 DRGs were cut on a cryostat and mounted on Superfrost Plus Microscope slides and air dried overnight at room temperature. Slides underwent pretreatment for fluorescence *in situ* hybridization consisting of 10 min Xylene bath, 4 min 100% ethanol bath, and 2 min RNAscope® H2O2 treatment. Next, the FISH protocol for RNAscope Fluorescent v2 Assay (ACD) was followed for hybridization to marker probes. Slides were then coverslipped with VECTASHIELD HardSet Antifade Mounting Medium with DAPI.

#### Microscopy

All images were captured on a Nikon Eclipse Ti2 microscope using a 20× or 40× objective and analyzed using NIS-Elements Advanced Research software v5.02. An examiner blinded to treatment and sex counted the number of positive pERK cells in laminae I-II. Cells with at least 3 puncta associated with a DAPI nucleus were considered positive for fluorescence *in situ* hybridization.

#### Statistical Analyses

All data are presented as means ± SEM. Statistical significance was determined as \**P* < 0.05. The effects of Drug and Time were analyzed by two-way analysis of variance (ANOVA), followed by Sidak’s multiple comparison tests. Data from dose-response curves were also analyzed as area under the curve using the trapezoidal method and used to produce the non-linear regression analyses of Maximum Possible Effect (% MPE). MPE were used to determine the ED50 for each drug. %MPE was calculated as follows: % MPE = 100 * (post-injection threshold – preinjection threshold)/(post-injury threshold – pre-injection threshold). All statistical analyses were performed in GraphPad Prism 9.0. Graphpad Prism and Biorender.com were used to make the graphics.

## Data Availability

All study data can be made available upon reasonable request.

## Author Contributions

T.S.N., D.F.S.S., and B.K.T. designed research; T.S.N, D.F.S.S., P.P., M.G., and H.N.A. performed research; T.S.N., D.F.S.S., P.P., and H.N.A. analyzed data; T.S.N. and B.K.T. wrote the manuscript with contribution from the other authors; T.S.N. and B.K.T. received funding for the project; B.K.T. supervised the overall project.

## Acknowledgments

This work was supported by National Institute of Health (NIH) grants R01DA37621, R01NS45954 and R01NS62306 to B.K.T and T32NS073548, F31NS117054, and F99NS124190 to T.S.N. We would also like to thank Drs. Herbert Herzog, Carmen Birchmeier, and Xinzhong Dong for providing key mouse lines that without which this work could not have been completed. We would like to thank Ronald Sivak for his essential assistance with genotyping the mouse lines. We thank Nina Gakii for critical experimental assistance throughout this project. We thank Dr. George Wilcox for assistance with the isobolographic analysis.

